# In Need of Age-Appropriate Cardiac Models: Impact of Cell Age on Extracellular Matrix Therapy Outcomes

**DOI:** 10.1101/2023.03.14.532565

**Authors:** S. Gulberk Ozcebe, Pinar Zorlutuna

## Abstract

Aging is the main risk factor for cardiovascular disease (CVD). As the world’s population ages rapidly and CVD rates rise, there is a growing need for physiologically relevant models of aging hearts to better understand cardiac aging. Translational research relies heavily on young animal models, however, these models correspond to early ages in human life, therefore cannot fully capture the pathophysiology of age-related CVD. Here, we chronologically aged human induced pluripotent stem cell-derived cardiomyocytes (iCMs) and compared in vitro iCM aging to native human cardiac tissue aging. We showed that 14-month-old advanced aged iCMs had an aging profile similar to the aged human heart and recapitulated age-related disease hallmarks. We then used aged iCMs to study the effect of cell age on the young extracellular matrix (ECM) therapy, an emerging approach for myocardial infarction (MI) treatment and prevention. Young ECM decreased oxidative stress, improved survival, and post-MI beating in aged iCMs. In the absence of stress, young ECM improved beating and reversed aging-associated expressions in 3-month-old iCMs while causing the opposite effect on 14-month-old iCMs. The same young ECM treatment surprisingly increased SASP and impaired beating in advanced aged iCMs. Overall, we showed that young ECM therapy had a positive effect on post-MI recovery, however, cell age was determinant in the treatment outcomes without any stress conditions. Therefore, “one-size-fits-all” approaches to ECM treatments fail, and cardiac tissue engineered models with age-matched human iCMs are valuable in translational basic research for determining the appropriate treatment, particularly for the elderly.

## 1. INTRODUCTION

Age is a significant risk factor for cardiovascular diseases (CVD), including myocardial infarction (MI). Studies show that more than half of CVD morbidity and long-term mortality following MI occur in individuals aged 65 years and older [1,2]. Age-related changes at the cellular, extracellular, and tissue levels negatively impact disease diagnosis as well as therapeutic outcomes. Pharmacological treatment outcomes were reported to be inconsistent and unpredictable for the elderly [3]. Similarly, despite the demonstrated benefits of cell-based therapies for MI in preclinical studies, early clinical trials resulted in limited improvement in left ventricular ejection fraction and ventricular remodeling, particularly for the elderly [4]. This is mainly due to decreased responsiveness of aged cells to their environment, and consequently to treatments [5,6]. Understanding cardiac aging and the effect that this has on CVD therapy outcomes are essential to ultimately prevent and treat age-associated disease syndromes.

Decellularized extracellular matrix (ECM) is a promising biomaterial for the regeneration and repair of musculoskeletal[7], neural[8], liver[9], and cardiovascular systems [10,11]. Studies have reported regenerative capabilities to be more effective when ECM was obtained from young tissues[6,12]. We previously showed the differences in the human induced pluripotent stem cell (iPSC)-derived cardiomyocytes (iCM) response to young, adult, and aged cardiac ECM. We showed that young ECM increases cell proliferation and drug responsiveness, improves cardiac function overall, initiates cell cycle re-entry, and mitigates oxidative stress damage in quiescent state aged iCMs [6]. Moreover, regardless of the ECM age, other groups have demonstrated the feasibility of using ECM for post-MI ventricular remodeling and cardiac functional recovery (i.e., LVEF) in animal models. Porcine cardiac ECM-derived hydrogels have been reported to increase the number of endogenous cardiomyocytes while preserving post-MI cardiac function[13]. Neonatal mouse cardiac ECM was shown to be more effective to prevent post-MI adverse ventricular remodeling, such as fibrosis, compared to adult ECM[14]. In another study, zebrafish heart ECM, which is known to be highly regenerative, was reported to exert pro-proliferative effects and contribute to post-MI cardiac regeneration in adult mice[15]. A recent clinical study showed that hydrogels derived from decellularized porcine myocardium improved left ventricular function in post-MI patients (57 to 62-year-old) [16]. Taken together, these studies provided valuable information on the safety, feasibility, and efficacy of ECM as a regenerative and post-MI therapy. There is a growing trend towards using ECM therapies, alone or in combination with cells, for MI. However, because current studies are largely based on young cells and young animal models, we still have a limited understanding of how the aged heart would respond to ECM therapies. As MI disproportionately affects the elderly, and the therapy outcomes vary with the patient age, here we generated an age-appropriate heart tissue model using aged iCMs and investigated ECM therapies for both MI treatment and prevention.

In this study, we explored age-related changes in the human heart. We characterized young (<30 years-old) and aged (>50-years-old) nonfailing human left ventricle (LV) samples and identified genes strongly altered with aging. We then compared the gene expression profiles of 3-month-old, 6-month-old, and 14-month-old (advanced aged) iCMs to those of human LV. Our results revealed a high degree of similarity between the advanced aged iCMs and the aged LV in terms of their stress and contractile function impairment related transcriptional signatures.

Next, we used chronologically aged iCMs to explore age-related changes *in vitro.* We investigated the effects of cell age on ECM therapy outcomes. Our findings showed that ECM treatment outcomes were influenced not only by cell age but also by the presence of stress conditions such as MI. Following MI-mimicking stress conditions (i.e., hypoxia), young ECM treatment led to functional recovery at all cell ages, with increased survival observed only in 3-month-old iCMs.

In the absence of stress, the beneficial effects of young ECM treatment were limited to the younger, 3-month-old iCM group. ECM upregulated cardiac structural and functional genes, increased beating frequency and velocity, and suppressed stress-related genes and the senescence-associated secretory phenotype (SASP) in 3-month-old iCMs. Surprisingly, young ECM was pro-aging and reduced the beating of advanced aged iCMs. These results challenged the widely held assumption of the universal benefit of young ECM, raising uncertainties about the safety and efficacy of employing young ECM as a preventative treatment for CVD in the elderly.

In conclusion, here we reported age-dependent transcriptional alterations in nonfailing human heart LVs from both young and aged subjects, and in chronologically aged iCMs. To the best of our knowledge, this is the only study displaying transcriptional alterations in human LV with a focus on aging without any disease conditions. Furthermore, our results showed that chronologically aged iCMs are excellent candidates to mimic aged heart behavior and can be used to conduct CVD studies for the elderly. Using an age-appropriate cardiac model, we showed that the ‘one-size-fits-all’ ECM treatment approach is doomed to fail, as results are highly dependent on cell age and stress conditions.

## 2. RESULTS

### 2.1. Distinct transcriptomic profiles differentiate aged human heart left ventricles from young counterparts

Left ventricles derived from healthy human hearts (young: <30-years-old, n=3 and aged: >50-years-old, n=3) were characterized. mRNA levels related to iCM maturity, function, and apoptosis were quantified and a 67% variance was detected between aged and young LV samples **(Fig.1A-B)**. Adult type sarcomeric genes (*TNNI3, MYH7*) and multiple Ca^2+^ cycling/SR genes were highly expressed in young LV **(Fig. 1C-D)**. Myocardial fibrillar collagens (*COL1A1, COL3A1*) along with other adverse cardiac remodeling contributors and stress-related genes were highly expressed in aged LV **(Fig. 1E-F).** Among screened genes, we identified 9 that were significantly altered (p<0.05) by human cardiac aging. Specifically, NPPB, a ventricular natriuretic peptide known to be secreted in the myocardium upon stress, was upregulated 40-fold (p=0.017) in the aged LV **(Fig. 1G)**. The KEGG pathway analysis revealed that the differentially expressed genes (DEGs) in young LV were associated with cardiac muscle contraction, calcium signaling, and focal adhesion pathways, while in aged LV they were associated with HIF-1, PI3K-Akt signaling pathways, hence cardiovascular aging, and cardiac disorders **(Fig. 1H)**. The results of gene ontology (GO) analysis also showed that DEGs in young LV were significantly enriched in biological processes, including the regulation of cardiac muscle contraction by calcium ion signaling, and cell communication. Aged LV DEGs were enriched in cGMP signaling pathways, cardiac muscle tissue development, and neuropeptide signaling pathways **(Fig. 1I)**. These changes in gene expression indicated that human cardiac aging mainly affected CM contractile function, Ca^2+^ cycling, and stress response.

**Figure 1.**
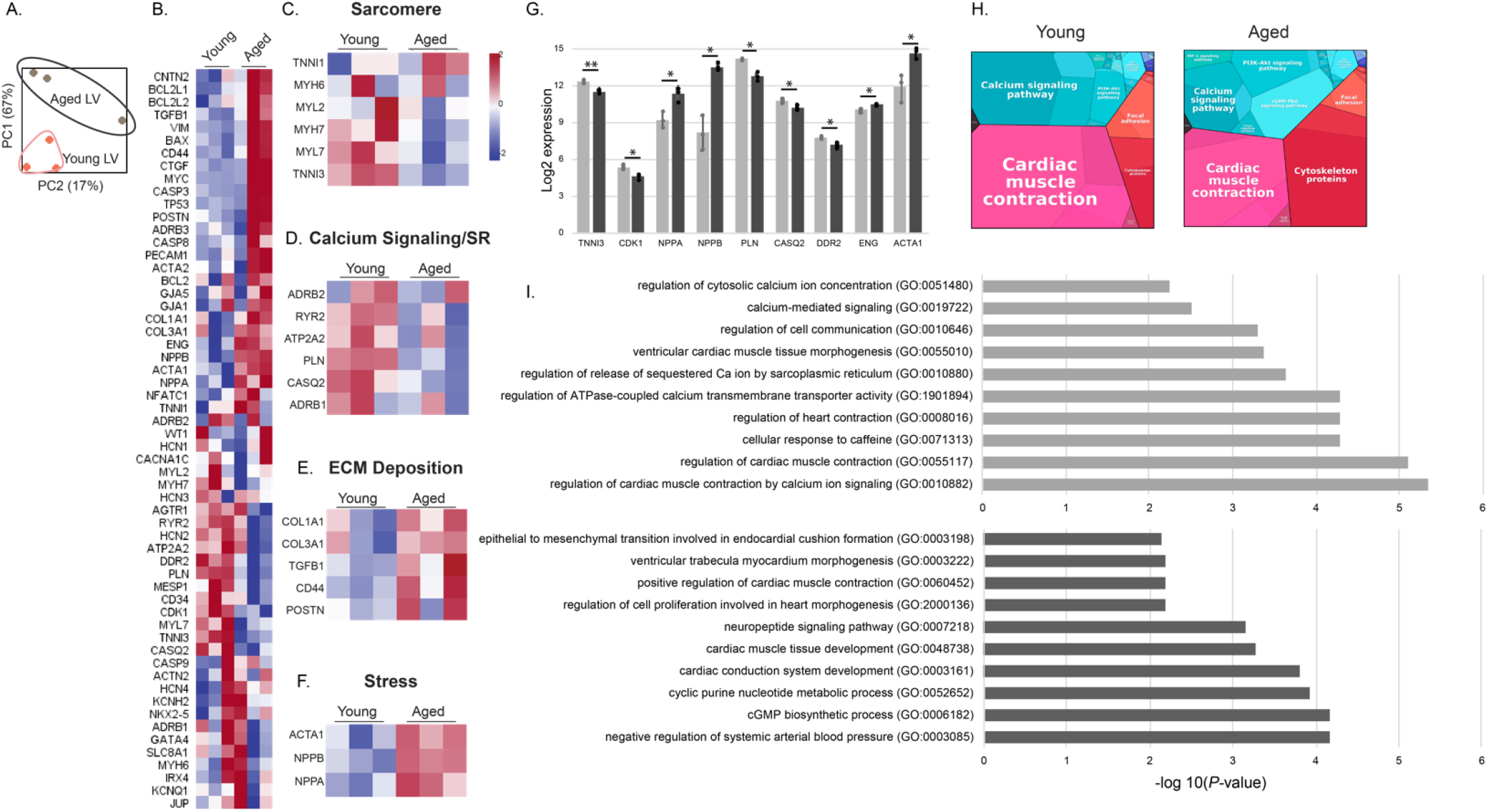
Human heart left ventricle age dependent transcriptional alterations. (A) Principal component analysis (PCA) of the gene expression data, depicting the group relationships of young (n=3) and aged (n=3) human left ventricles (LV). The proportion of component variance is indicated as a percentage. (B) Differential expression levels of the preselected 58 cardiac and aging specific genes for young and aged LVs. (C-F) Heatmaps of key genes involved in distinct features of CM behavior: (C) sarcomere, (D) calcium (Ca2+) cycling and sarcoplasmic reticulum (SR), (E) ECM deposition, and (F) stress response. (G) Statistically altered gene expressions of young and aged human LV. Statistical analysis was done using one-way ANOVA with post-hoc Tukey’s test. **p<0.01, *p<0.05, n≥3. Data presented as mean ± standard deviation (SD). (H) Proteomaps showing the KEGG pathways. (I) Gene ontology analysis showing the biological processes associated with the genes overexpressed in young and aged LVs.

### 2.2. Transcriptomic alterations in chronologically aged iCMs

Unsupervised hierarchical clustering revealed a 66% variance between young and aged iCMs **(Fig. 2A-B)**. GO analysis revealed that genes downregulated in advanced iCMs were significantly enriched in biological processes, including DNA damage response, apoptotic processes, and cell cycle progression. The genes that were upregulated with iCM aging were associated with cardiac contraction, ion transport, and calcium signaling **(Supp. Fig. 1)**, indicating acquired structural and functional maturity with prolonged culture time. With prolonged culture, the adult type sarcomeric myosins and cardiac troponin were differentially expressed **(Fig. 2C, H)**. The intermediate filament protein (*VIM*) that is highly expressed in fetal CMs was downregulated with cellular aging. Although structurally more mature, advanced aged iCMs grouped together with young 1-month-old iCMs regarding their calcium signaling and sarcomeric reticulum related expressions **(Fig. 2D)**. Reduced excitation-contraction coupling expressions were observed in both immature 1-month-old and advanced aged iCMs, indicating that CMs reverted toward an impaired calcium handling machinery as they age. The calcium handling genes that are essential for cardiac action potential and cardiac contraction were upregulated up to 8-fold in 3- and 6-month-old iCMs, while the negative calcium import regulator (*PLN*) was upregulated in 6- and 14-month-old iCMs. Consequently, *PLN:ATP2A2* ratio, an indicator of reduced SERCA activity and impaired calcium handling, was more than doubled in 6- and 14-month-old iCMs **(Supp. Fig.1C)**. Potassium and sodium channel gene expressions were gradually increased with prolonged culture **(Fig. 2E, H)**.

**Figure 2.**
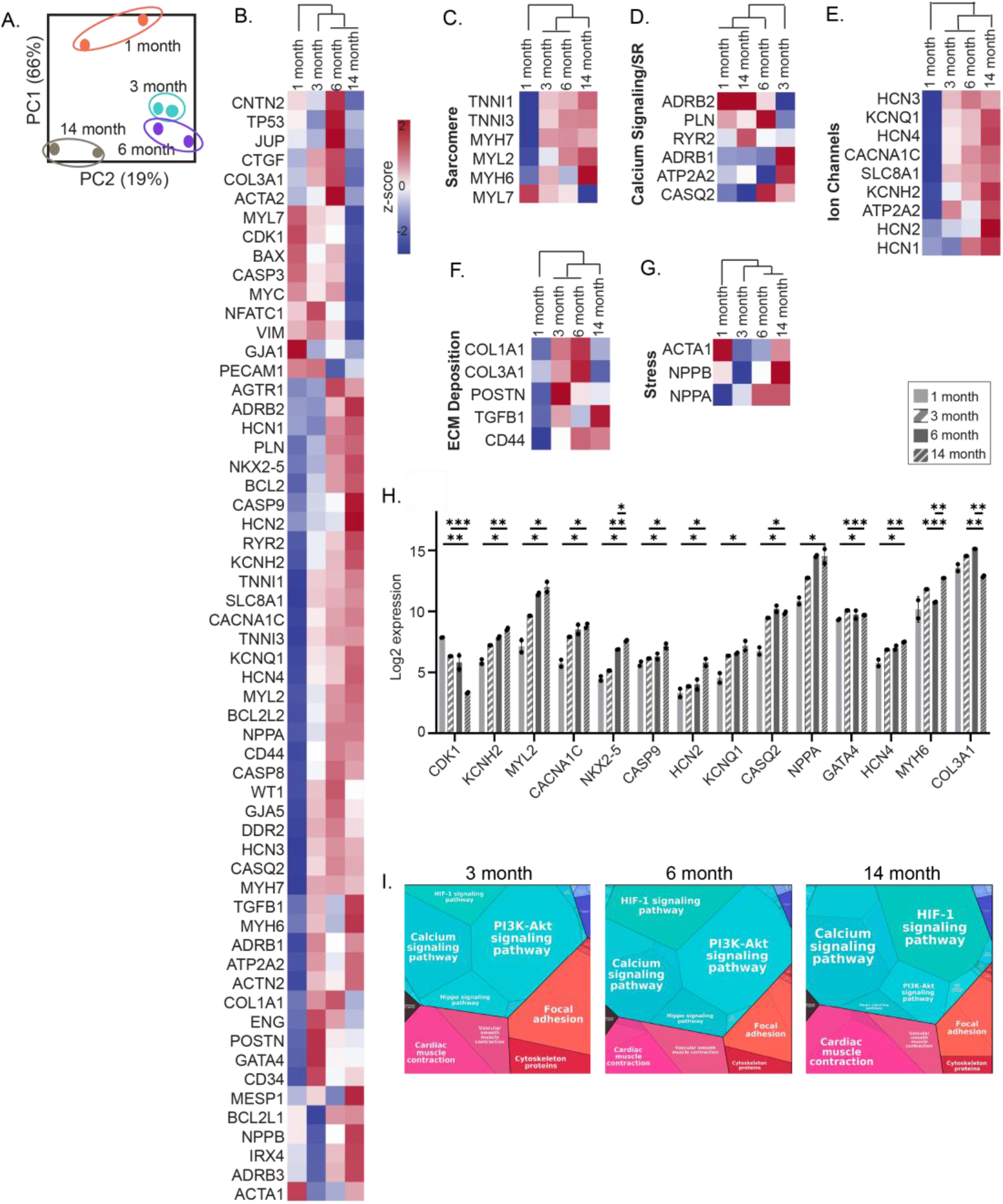
iCM age-dependent transcriptional alterations. (A) Principal component analysis (PCA) of the gene expression data, depicting the group relationships of young and aged iCMs. (B) Differential expression levels of the preselected 58 cardiac and aging specific genes for iCMs. (C–G) Heatmaps of key genes involved in distinct features of CM behavior: (C) sarcomere, (D) calcium (Ca2+) cycling and sarcoplasmic reticulum (SR), (E) ion channels, (F) ECM deposition, and (G) stress response. (H) Statistically altered gene expressions of aged iCMs. Statistical analysis was done using one-way ANOVA with post-hoc Tukey’s test. ***<0.001, **p<0.01, *p<0.05, n=6 pooled into 2 technical replicates. Data presented as mean ± standard deviation (SD). (I) Proteomaps showing the KEGG pathways.

In agreement with the human LV gene profile, adverse cardiac remodeling genes that mediate cardiac fibrosis were upregulated with iCM aging **(Fig. 2F)**. As in aged LV, we detected high levels of cardiac hypertrophy and stress related (*NPPA, NPPB*) expressions in advanced aged cells. Especially *NPPA* was upregulated more than 16-fold from 1-month to 14-month of culture **(Fig. 2G)**. Although experiencing more stress, and significantly upregulated initiator caspase *CASP9* expression **(Fig. 2H)**, a predictive value that determines the susceptibility to the apoptotic signal, *BAX/BCL2* ratio revealed that advanced aged cells were 2 times more resistant to apoptosis at the gene level **(Supp. Fig.1 B)**.

**Supplementary Figure 1.**
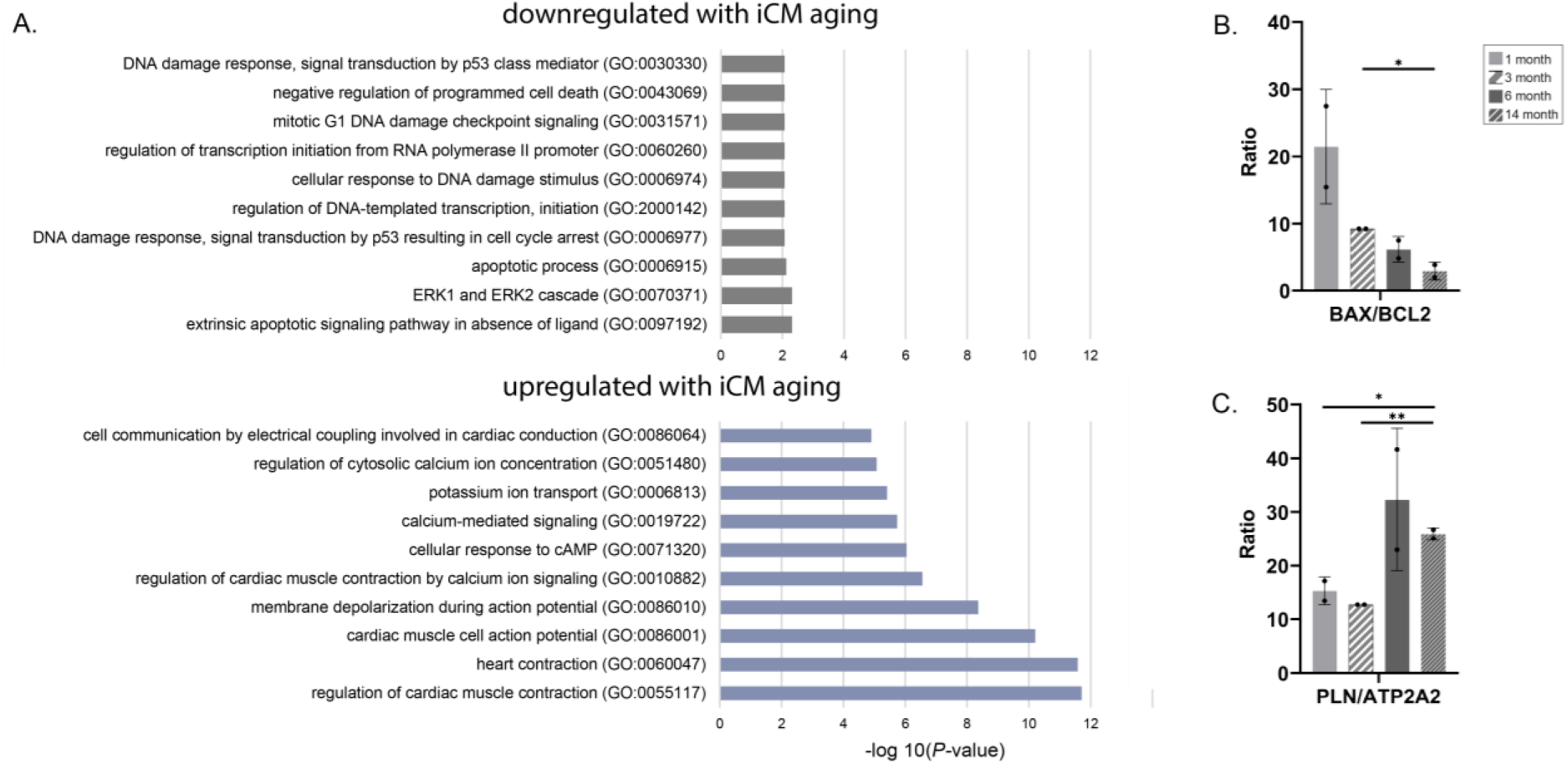
iCM age-dependent DEG alterations. (A)Gene ontology analysis showing the biological processes associated with the genes down- and up-regulated with chronological aging of iCMs. Calculated (B) BAX/BCL2 and (C) PLN/ATP2A2 ratios. Statistical analysis was done using one-way ANOVA with post-hoc Tukey’s test. ***<0.001, **p<0.01, *p<0.05, n=6 pooled into 2 technical replicates. Data presented as mean ± standard deviation (SD).

Additionally, cell-cycle-associated *CDK-1* gene expression was significantly downregulated while *GATA4, a* critical regulator of cardiac regeneration, and its cofactor, *NKX2-5* expressions were upregulated in advanced aged iCM **(Fig. 2H)**. The KEGG pathway analysis of the relative expression data further showed that regeneration mediator Hippo signaling and PI3K-Akt pathways were downregulated, while HIF-1 signaling pathway and cardiac muscle contraction were upregulated in advanced aged iCMs **(Fig. 2I).**

Advanced aged iCMs had a larger proteome body, which might indicate increased age-associated inflammation or inflammageing. Among the highly detected proteins, the senescence mediator (SERPINE1) increased whereas the critical mediator of cardiovascular health (ENG) decreased gradually with cellular aging **(Fig. 3A-B)**. In agreement with the transcriptomic alterations, the pro-aging Ras signaling pathway, master senescence associated secretory phenotype (SASP)-regulator NF-KB signaling pathway, HIF1α and cytokine-cytokine receptor interaction proteins that are known to be associated with age-related degenerations in multiple organ systems[17,18] were highly expressed in advanced aged iCMs **(Fig. 3B)**.

**Figure 3.**
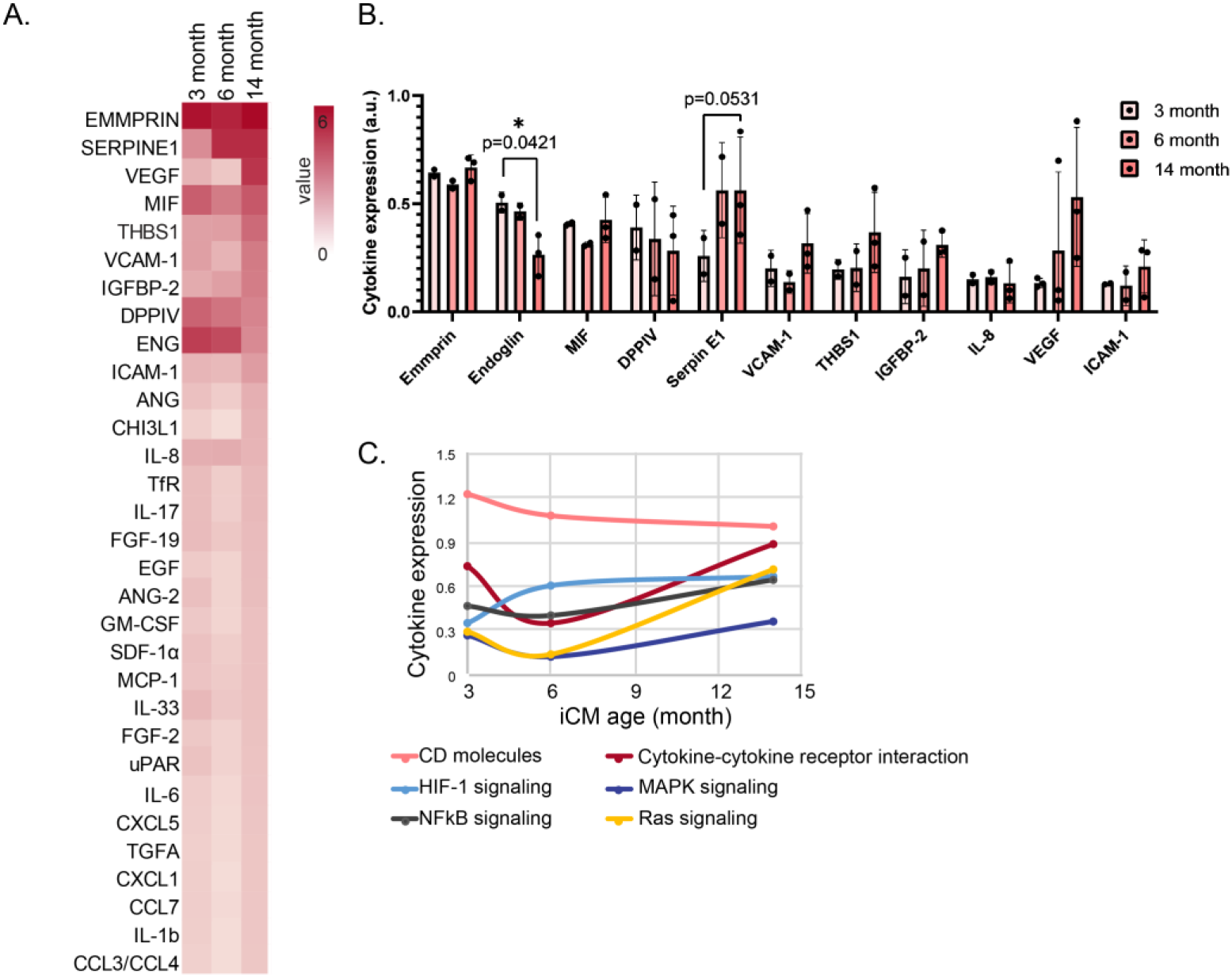
Aged iCM cytokine expression. (A) Heatmap showing the screened protein expressions of aged iCMs (B) Highly expressed cytokine expressions of aged iCMs. Statistical analysis was done using one-way ANOVA with post-hoc Tukey’s test. ***<0.001, **p<0.01, *p<0.05, n=6 pooled into ≥2 technical replicates. Data presented as mean ± standard deviation (SD). (C) iCM age dependent changes in the cytokine expressions with respect to their role or involved pathways.

### 2.3. ECM treatment effect on post-MI functional recovery

To determine whether there were any cell age-dependent responses to the young ECM treatment for post-MI recovery, we exposed 3-month-old and advanced aged iCMs to MI-like stress conditions. The spontaneous beating of the cells was recorded before anoxia, and at 1h, 6h, and 12h RI to assess the beating recovery, and beating frequency **(Fig. 4A)**, beating velocity **(Fig. 4B)**, maximum displacement **(Fig. 4C)**, and beating area **(Fig. 4D)**. 3-month-old iCMs had higher initial beating frequencies (0.59±0.04 Hz) than the advanced aged iCMs (0.22±0.02 Hz), hence beating values were normalized separately.

**Figure 4.**
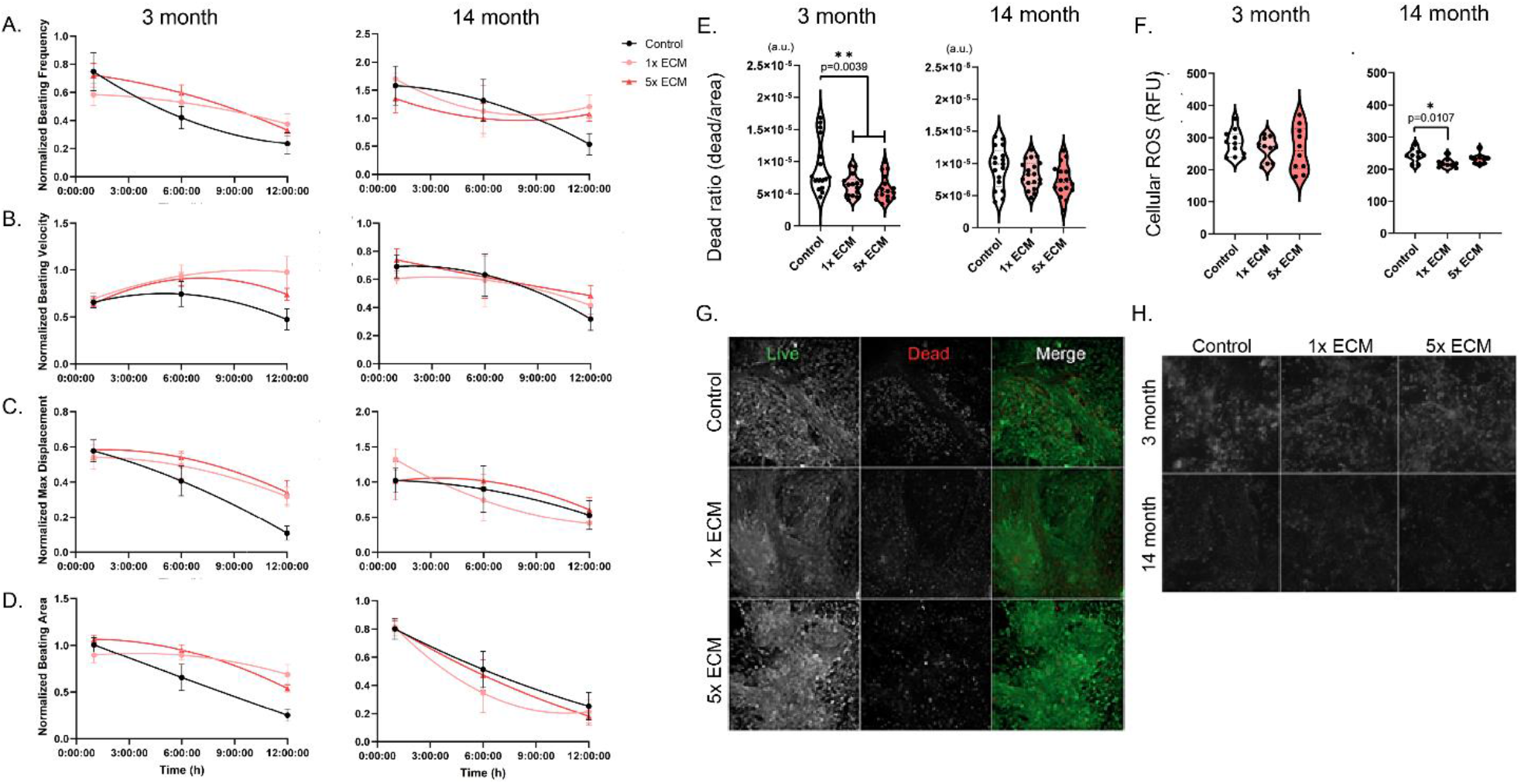
Post MI analysis. Temporal changes of (A) beating frequency, (B) beating velocity of spontaneous cell beating, (C) maximum displacement of a pixel in a frame due to spontaneous beating, and (D) beat area (%) recorded for 12h post-MI. Left panel:3-month-old, right panel:Advanced aged iCM. Post-MI (E) cell death ratio and (F) cellular ROS levels of 3-month-old 14-month-old advanced aged iCMs at 12h RI. Representative images of (G) 3-month-old iCM live dead staining and H) ROS generation measurement of 3-month-old and advanced aged iCM at 3h RI. Statistical analysis was done using one-way ANOVA with post-hoc Tukey’s test. **p<0.01, *p<0.05, n≥3. Beating recovery was normalized to the pre-MI initial values and data presented as mean ± standard deviation (SD).

Within the first hour post-anoxia, 3-month-iCMs slowed down while advanced aged iCMs displayed rapid, irregular twitches resulting in doubled beating frequencies. At the end of 12h, 1-out-of-4 samples of 3-month-old, and 4-out-of-10 samples of advanced aged iCMs stopped beating. We observed beneficial effects of ECM treatment in both 3-month and advanced aged iCMs. Control untreated 3-month-old group beating frequency, robustness, and area recovered to only half of their original pre-MI values, while ECM treated groups had faster, stronger beating across a larger area at the end of 12h normoxia **(Fig. 4A-D)**. For advanced aged iCMs, ECM treatment sustained their beating frequency **(Fig. 4A),** and the control group frequency decreased to half of the original value. However, ECM treatment did not affect the beating robustness or area. Regardless of the ECM, the beating area of advanced aged iCMs was dramatically reduced.

### 2.4. ECM treatment effect on the deleterious effects of MI

We investigated the effect of cell age on the therapeutic potential of the young ECM. Regardless of the dose used, ECM significantly lowered the dead cell count of the 3-month-old cells (p=0.0039) while the detected cellular ROS levels were comparable **(Fig. 4E-H)**. There was no survival difference in the advanced aged iCMs **(Fig. 4E)**, however, mitochondrial ROS generation significantly decreased (p=0.0107) with the ECM supplementation **(Fig. 4F)**. Advanced aged cells were further screened for apoptosis-related proteins to investigate why we detected ECM effects on ROS generation but not on cell survival. When relative expressions were compared, we detected high levels of clusterin, cell protectant protein against ROS-induced apoptosis, ROS scavenger catalase, and low levels of stress proteins HSP60 and HSP70 in ECM supplemented cells. The pro-apoptotic proteins did not show a difference, yet one of the critical early mediators of apoptosis, Ser46 phosphorylation of p53 level, and the apoptosis initiator, Cytochrome C gradually decreased with increasing ECM dose (**Supp. Fig. 2)**.

**Supplementary Figure 2.**
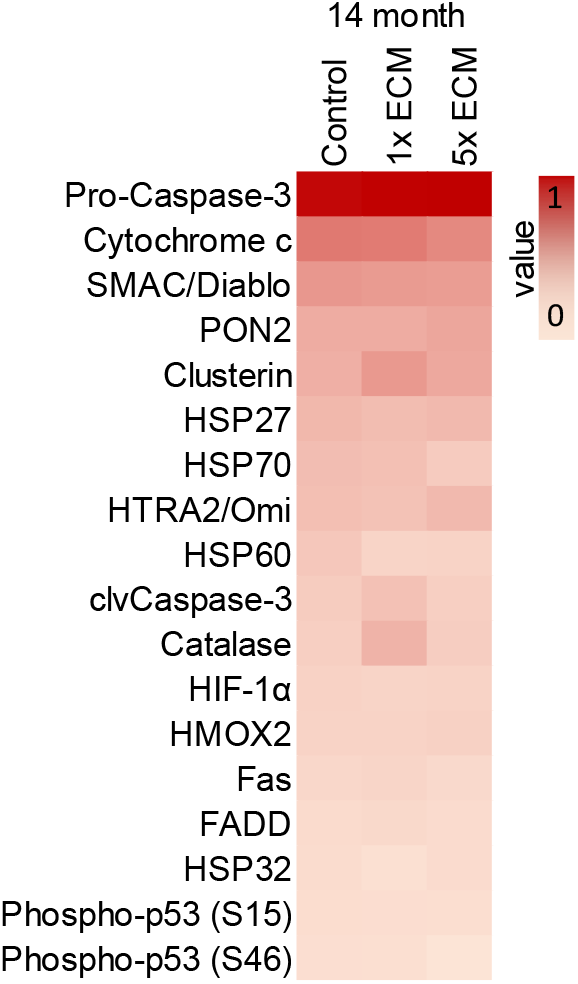
Post-MI apoptosis-related protein expression in 14-month-old advanced aged iCMs. n≥3.

### 2.5. ECM treatment effect on aged iCM expressions without any stress conditions

Following a 10-day treatment with young human heart LV-derived ECM at two concentrations (1x ECM: 0.1mg/ml and 5x ECM: 0.5mg/ml), we investigated the changes in transcriptome and cytokine levels. Regardless of the cell age or the ECM concentration, the treatment downregulated genes that are associated with collagen activated signaling pathways, extracellular organization, and non-cardiac cell migration, and upregulated genes associated with cardiac cell fate commitment and muscle contraction **(Fig. 5A)**. GO analysis also showed that genes upregulated upon ECM treatment were enriched in regulation of glial cell differentiation, which we recently reported to be regulators of heart rate [19].

**Figure 5.**
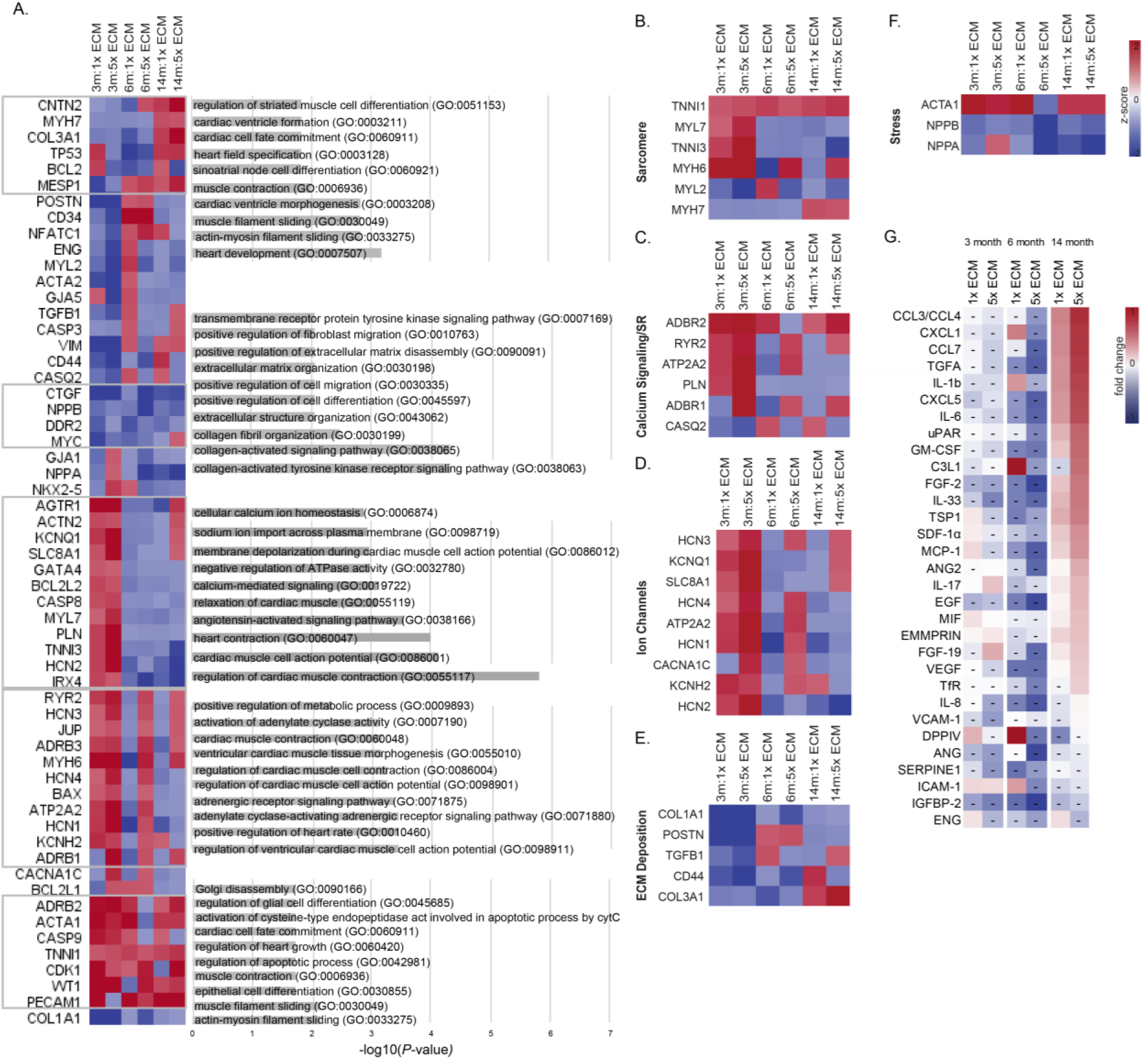
ECM treatment effect on aged iCMs. (A) Differential expression levels of the preselected 58 cardiac and aging specific genes for iCMs. Gene ontology analysis showing the biological processes associated with the up- and downregulated genes with iCM aging. (B–F) Heatmaps of key genes involved in distinct features of CM behavior: (B) sarcomere, (C) calcium (Ca2+) cycling and sarcoplasmic reticulum (SR), (D) ion channels, (E) ECM deposition, and (F) stress response. (G) Heatmap showing the screened protein expressions as a fold change from their corresponding control groups. n≥3.

Expectedly, high concentration ECM had a greater effect on the cells, and the ECM treatment outcome depended heavily on the cell age. GO results showed that DEGs in 3-month-old iCMs were associated with functional processes such as the movement of ions and cardiac muscle contraction and relaxation. Relatedly, sarcomere, calcium signaling, and ion channel expressions were upregulated in an ECM dose dependent manner in 3-month-old iCMs **(Fig 5B-D)**. DEGs in 3-month-old and high dose ECM treated 6- and 14-month-old iCMs were involved in the positive regulation of metabolic processes in addition to cardiac contraction, and regulation of cell action potential. Adrenergic receptor (*ADRB1*) and ryanodine receptor (*RYR2*) levels which play an integral roles in excitation-contraction coupling and cardiac energy metabolism **(Fig. 5C)**, increased with high concentration ECM treatment. HCN channel family, Ca^2+^ and K^+^ channel levels in 6-month-old iCMs, and Na^+^/Ca^2+^ exchanger (*SLC8A1*) levels in advanced aged iCMs increased only when treated with high concentration ECM. Surprisingly, DEGs in 14-month-old iCMs treated with ECM were associated with heart development and cardiac ventricle morphogenesis **(Fig.5A)**. Besides cardiac structure and function, ECM treatment downregulated ECM deposition genes, especially in 3-month-old iCMs. *COL3A1* was upregulated only in advanced age iCMs in an ECM dose-dependent manner **(Fig. 5E).** Regarding cellular stress, the genes we have dominantly seen in the advanced aged iCMs (*NPPA, NPPB*) were downregulated in all and were almost halved with the high concentration ECM treatment **(Fig. 5F)**.

At the protein level, ECM lowered the SASP components increased with cellular aging, such as the aging markers SERPINE1, IL-8, IGFBP-2, and VCAM **(Fig. 5G)**. ECM also decreased the maladaptive aging response associated cytokine, and chemokines (THBS1, CHI3L1, IL33) in 3-month and 6-month-old iCMs. However, ECM treatment had a different effect on the advanced aged iCM than on 3-month and 6-month-old iCMs **(Fig. 5G)**. Pro-aging Ras and MAPK signaling pathway proteins (i.e., IL-1b, FGF2) were highly expressed in advanced aged iCMs.

### 2.6. ECM treatment effect on aged iCM beating without any stress conditions

We recorded the spontaneous beating of the aged iCMs on day 3 and day 10 of the ECM treatment. The initial beating frequencies were recorded as 0.72 ±0.24 Hz for 3-month, 0.48 ±0.19 Hz for 6-month and 0.25 ±0.10 Hz for advanced aged iCMs, expectedly decreasing with increasing cell age. ECM treatment enhanced the beating of 3-month-old iCMs, whose cells reached adult CM beating frequency (~1 Hz) and had significantly increased beating velocities at the end of 10-day high dose ECM treatment **(Fig. 6A-B)**. However, we observed minimal if not negative effects of ECM treatment on the 6-month-old and advanced age iCM beating properties **(Fig. 6A-C)**.

**Figure 6.**
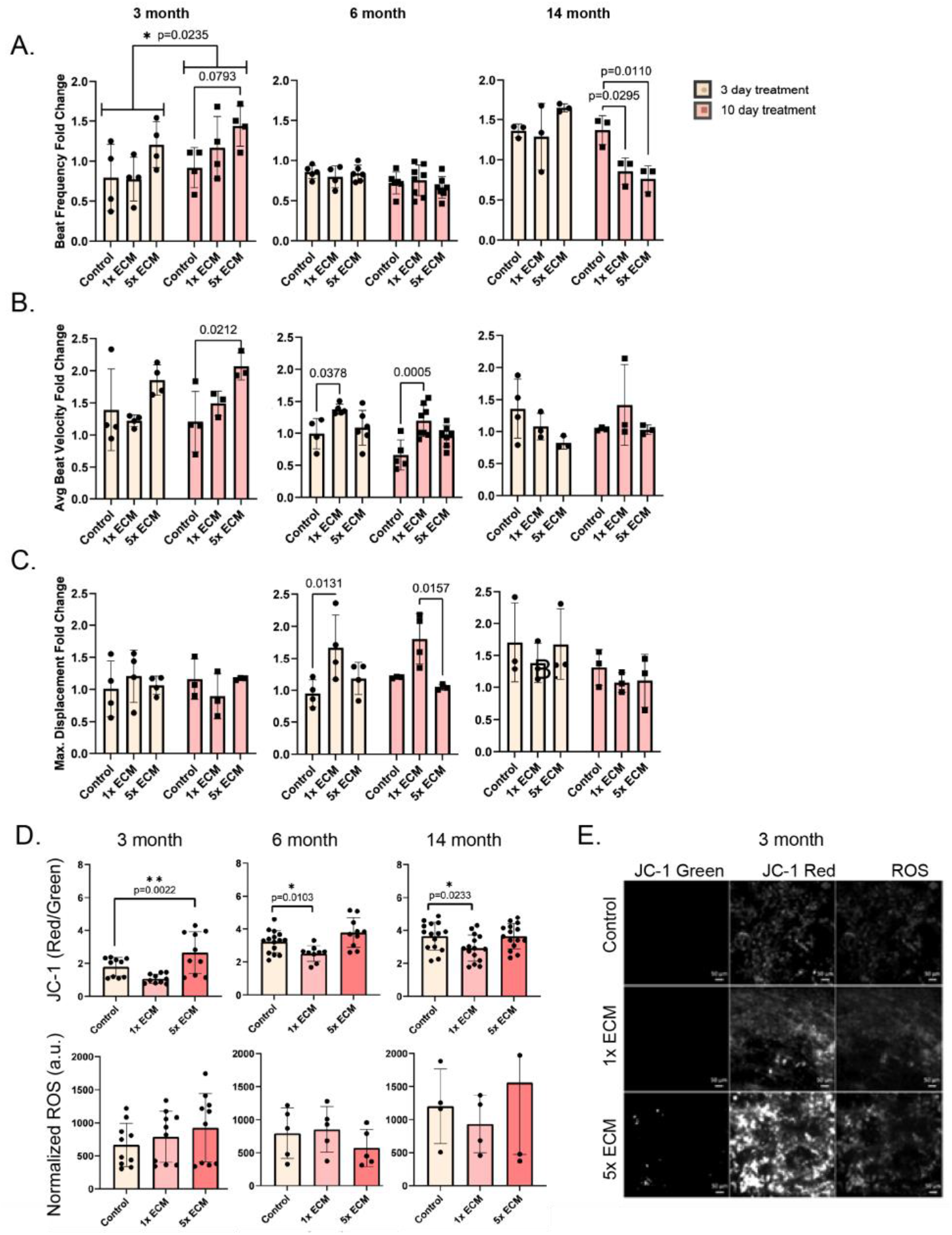
ECM treatment effect on aged iCM beating kinetics. Fold change of (A) beating frequency, (B) beating velocity of spontaneous cell beating and (C) maximum displacement of a pixel in a frame due to spontaneous beating on day 3 and 10 of treatment. (D) Quantification of mitochondrial potential (JC-1 Red/Green) as an indication of the mitochondrial health (top) and ROS levels normalized to pre-treatment measurement (bottom). Left panel: 3-month, middle-panel:6-month, right panel:advanced aged iCM. (E) 3-month-old iCM mitochondrial staining with JC-1 dye and cellular staining with ROS on the 10th day of ECM treatment. Scale bar: 50μm. Statistical analysis was done using one-way ANOVA with post-hoc Tukey’s test. **p<0.01, *p<0.05, n≥3. Data presented as mean ± standard deviation (SD).

Relatedly, mitochondrial health was assessed, and we detected higher mitochondrial membrane potential (Red/Green ratio), indicating increased ATP generation potential in 3-month-old iCMs after 10-day ECM treatment **(Fig. 6D)**. Interestingly, the mitochondrial potential for both 6-month and advanced aged iCMs significantly decreased after 1x ECM treatment **(Fig. 6D)**. Referring to the strong correlation between mitochondrial membrane potential and ROS production, we also measured the cellular ROS levels and observed only minimal changes with cell age or ECM treatment. Although ECM increased mitochondrial activity in 3-month-old iCMs, it did not lead to increased ROS production **(Fig. 6E)**.

## 3. DISCUSSION

In this study, we compared aging profiles of human LV and chronologically aged iCMs, and developed aged human heart models using aged iCMs to investigate the effect of cell age on young ECM treatment outcomes. We demonstrated that young ECM is an effective treatment for MI, as it promoted the survival of cells and facilitated their beating recovery, particularly in younger iCMs. However, young ECM had unexpected negative effects as a preventative therapy for advanced aged iCMs in the absence of any stress conditions. Despite its origin in young human LV, ECM increased aging-factor expression and impaired beating of advanced aged iCMs, challenging its use for preventative purposes, especially for the elderly.

Many researchers have presented the human LV transcriptome in association with various diseases and conditions, yet we don’t have a complete understanding of human heart aging. Age related changes reported for rodent hearts include myocyte hypertrophy, cardiac fibrosis, and reduced calcium transport across the sarcoplasmic reticulum membrane [21]. Here, we have confirmed the previously described age-related pathophysiology at the transcriptomic level in human heart left ventricles (LV). We observed elevated levels of NPPA and NPPB in all aged LV samples **(Fig.1)**, which is considered a hallmark of human aging and a protective hormonal response to mechanical stress to maintain cardiovascular homeostasis [22]. Additionally, DEGs in aged LVs showed an association with cGMP and HIF-1 signaling pathways, as well as negative regulation of JUN kinase activity leading to hypertrophy and pronounced fibrosis in aged LVs [23].

Prolonged culture (>1-month) has typically been used as a CM maturation strategy[24–27]. Therefore, little was known about the capacity of aged iCMs to mimic human cardiac aging. Here, we report for the first time that functional (i.e., beating) 14-month-old advanced aged iCMs exhibit the hallmarks of cardiac aging at both transcriptional and translational levels, including adverse cardiac modeling, hypertrophy, and SASP **(Fig. 2-3)**. We observed re-expression of fetal genes (i.e., *TNNI1* and *MYH6*), induction of pre-ANF (*NPPA*), pre-BNP (*NPPB*), and *ACTA1* in the advanced aged iCMs similar to the distinct molecular phenotype associated with pressure overload-induced hypertrophy[28]. Additionally, advanced aged iCMs displayed beta-adrenergic receptor expression levels that resemble those of 1-month-old iCMs **(Fig. 2D-E)**. The predominant expression of *ADRB2* (fetal-type) and low expression of *ADRB1* (adult-type) indicate β-adrenergic desensitization can be seen during cardiac aging as well as in immature CMs. Early studies suggest that the heart develops immature features with aging, including a dependence on transsarcolemmal calcium influx during contraction rather than calcium stored in the sarcoplasmic reticulum as in the adult heart [29,30]. Here, we showed that 14-month-old advanced aged iCMs were indeed senescent and highly similar to the aged LV at the transcriptional level.

We observed another phenomenon commonly seen in senescent cells. Despite high levels of initiator caspase CASP9, advanced aged iCMs had low levels of CASP3 **(Fig. 2B, H)**, indicating the central apoptotic machinery downstream CASP9 was inactivated and the upregulation of CASP9 was independent of apoptosis. Additionally, the increased resistance of advanced aged iCMs to apoptosis **(Supp. Fig.1B)** suggests a survival strategy specific to advanced aged iCMs, as also observed in aged olfactory bulb neurons but not in young counterparts [31]. This is known as the trade-off between senescence and apoptosis [32], and the surprising results of ECM treatment in the absence of stress conditions might be due to this delicate balance. Using the heart tissue model with different aged iCMs, we demonstrated the critical role of ‘cell age’ in determining ECM treatment efficacy and outcome. Consistent with previous preclinical and phase I clinical studies [6,13–16], young ECM improved post-MI beating recovery**(Fig. 4A-D)**. However, the effect of ECM on the post-MI survival rate was highly cell age dependent, with only the younger cell group showing higher survival in response to ECM treatment **(Fig.4E-H)**. Although young ECM enhanced oxidative stress coping mechanisms in advanced aged iCMs **(Supp. Fig. 2)**, a similar effect shown in a recent study revealing the ROS scavenger activity of ECM [33], this did not translate into an increase in their survival rate. The desensitization of advanced aged cells due to reduced cellular activity and function has long been known and also observed in the human heart as it ages [6,34]. However, such a difference in previous studies has not been reported because samples are pooled together regardless of age to show the global effect of ECM therapies.

Current MI guidelines do not differentiate treatment based on age or sex, hence the treatment efficacies are suboptimal in the elderly and women[35]. For this study, we acknowledge the potential sex-based differences at both the gene and protein levels. A recent study on transcriptional diversity of the human heart (n=7, ages: 39-60) reported 17 genes that exhibited sex-based differential expression within cardiomyocytes (i.e., *NEB, PBX3*) [36]. Another study on sex-related protein expressions in hypertrophic cardiomyopathy patients (n=26, ages: 48.5 ± 17.7 (F) and 49.8 ± 15.5 (M)) reported 46 proteins that were differentially expressed in the female and male groups (i.e., tubulins and HSPs)[37]. However, since we found no evidence for a sex-based differential expression in the genes or proteins of interest **(Fig. 1-3)**, we didn’t separate our samples by sex.

Studies have demonstrated the great potential of young ECM therapies in promoting post-MI recovery and regeneration, suggesting its use for preventative purposes. We demonstrated similar transcriptomic and related translational changes reported for post-MI ECM therapy results including reduced CM apoptosis, improved function, and cardiac development for 3-month-old iCMs treated with ECM **(Fig. 5A, 6)**. In addition, ECM treatment improved structural and functional cardiac maturity and decreased SASP (i.e., SERPINE1, IGFBP2, and interleukins), ECM deposition, and stress-related expressions in 3-month-old iCMs in a dose dependent manner **(Fig. 5B-F)**. We acknowledge the immature nature of iCMs, therefore observed maturation with the ECM treatment was expected. However, the ECM effect on SASP is noteworthy as SASP-centered approaches are emerging as alternatives to target senescence-associated diseases.

When we repeated the same ECM treatment for the 14-month-old advanced aged iCMs, we got unexpected results. The impact of ‘cell age’ on the outcome of ECM treatment was particularly significant in the absence of stress conditions. In fact, advanced aged iCMs were minimally or negatively affected by the ECM treatment. ECM increased SASP, namely CXC chemokines and activated IL-6/JAK-STAT pathway **(Fig. 5G)** in advanced aged iCMs suggesting that young ECM exerted hypertrophic stress on advanced aged iCMs. As per our observations on the beating properties of advanced-aged iCMs **(Fig. 6A)**, the elevation of pro-inflammatory cytokines is often associated with impaired cardiac function [38]. Moreover, a highly conserved pro-aging RAS/MAPK signaling pathway was upregulated in advanced aged iCMs after ECM treatment in a dose dependent manner **(Fig. 5G)**.

Old age is associated with worse treatment outcomes and patient age is determinant in decision-making and treatment selection in many disease conditions, including, breast cancer[39], and schizophrenia[40]. This study highlights that age is also a critical determinant in the treatment of CVDs. Despite recent advances in ECM therapies, its efficacy and outcomes in elderly patients remain limited by the lack of data. Our results clearly demonstrated that the advanced aged iCMs, representing the elderly, did not benefit equally from the post-MI young ECM treatment as the younger iCMs, and were even adversely affected in the absence of stress conditions. Therefore, age-appropriate cardiac models, such as the one presented here, are needed in the cardiac tissue engineering field to facilitate CVD therapy studies and enhance our understanding of cardiac aging.

## 4. CONCLUSION

Our study revealed age-dependent transcriptional alterations in nonfailing human heart LVs, with a sole focus on aging without any co-existing disease states. Moreover, we showed that chronologically aged iCMs are excellent candidates to mimic aged heart behavior, and aged heart models using age-appropriate iCMs are valuable for studying age-dependent efficacy and outcome of the CVD therapies. Our results demonstrated that the ECM response is highly dependent on cell age and stress conditions. Therefore, there is a need for age-appropriate cardiac models in translational research to develop personalized treatments for the elderly population, and to move beyond the “one-size-fits-all” approach in ECM therapies.

## 5. MATERIALS AND METHODS

### 5.1. Donor heart harvest

De-identified human hearts that were deemed unsuitable for transplantation and donated to research, were acquired from Indiana Donor Network under the Institutional Review Board (IRB) approval for deceased donor tissue recovery. Human heart tissues were grouped as young (from <30 years-old patients, n=3), and aged (from 50< years-old patients, n=3). For storage, hearts were dissected into its chambers and kept separately in a −80°C freezer until use. We only used the young left ventricles (n=3) for the ECM treatments

### 5.2. Decellularization of human heart tissue for matrix preparation

Left ventricles from young donors were sectioned and decellularized following previous decellularization protocol [41]. Briefly, we first stripped the fatty tissue around the left ventricular myocardial tissue and sliced the tissues in thin sections (<1mm). To decellularize, tissues were washed in 1% (wt/vol) sodium dodecyl sulfate (SDS) (VWR, #97062) for 24 hours or until white transparent tissue was obtained, then in 1% (wt/vol) Triton 100-X (Sigma-Aldrich, #A16046) for 30 minutes. After decellularization, samples were washed thoroughly with DI water to remove any residual detergent. To delipidize, tissues were washed with the isopropanol (IPA) for 3 hours then rehydrated in DI and treated with 50U/ml DNase (Millipore Sigma, #10104159001) for 8 hours followed by an overnight DI rinse. All steps were conducted with constant agitation at RT.

Prepared ECMs were lyophilized and pulverized with liquid nitrogen. ECM powder was digested in a 1 mg/mL pepsin (Sigma-Aldrich, #P6887) in 0.1M HCl (10:1, w/w, dry ECM:pepsin) at RT with constant stirring until a homogeneous solution was obtained. The insoluble remnants were removed by centrifugation, the supernatant was neutralized using 1M NaOH solution, and used immediately to prevent degradation. Prior to experiments, we measured the total protein concentrations using Rapid Gold BCA Assay (Thermo Scientific, # A53227) and diluted ECM solutions to either 0.01 mg/ml (1x) or 0.05 mg/ml (5x) with the culture media.

### 5.3. Human iPSC cell line

The cell line used in this study is DiPS 1016 SevA (RRID: CVCL_UK18) from human dermal fibroblasts obtained from Harvard Stem Cell Institute iPS Core Facility. Cells were cultured in humidified incubators at 37 °C and 5% CO2. Human iPS cells were cultured routinely in mTeSR-1 media (StemCell Technologies, #05825) on 1% Geltrex-coated plates (Invitrogen, #A1413201). At 80-85% confluency, cells were passaged using Accutase (StemCell Technologies, #07920) and seeded at 1.5 × 10^5^ cells/cm^2^ on well plates with Y-27632 (ROCK inhibitor, 5*μ*M), (StemCell Technologies, #129830-38-2) in mTeSR-1 media. The culture was maintained with daily media changes until 90% confluency was reached.

### 5.4. Culturing iPSC-derived cardiomyocytes

Once 90% confluency was reached, cardiac differentiation was initiated following canonical Wnt pathway [42]. To direct cardiac differentiation, cells are sequentially treated with CHIR99021 (12*μ*M) (Stemcell Technologies, #72052) for 24 hours followed by RPMI 1640 medium with B-27 supplement without insulin (2%) (Gibco, #A1895601) (CM(-)). Cells were then treated with Wnt pathway inhibitor IWP-4 (5 μM) (Stemcell Technologies, #72552) for 48 hours followed by CM(-) for 48 hours. From day 9 on, cells were maintained in RPMI 1640 medium with B-27 (2%) (Gibco, #17504044) (CM(+)) and media was changed every 3 days. iCMs were cultured for 3-months, 5-6-months and 13-14-months.

### 5.5. ECM treatment experiments

Myocardial infarction (MI) experiment was mimicked in two parts as ischemic phase (I) and reperfusion injury (RI). Aged cells were incubated under anoxic conditions (37 °C, 5% CO_2_, 0.1% O_2_) for 3 hours (I), then moved to normoxic conditions (21% O_2_) for 12 hours (RI). During ischemia, cells were incubated in anoxia-equilibrated RPMI 1640 medium without glucose (Corning, #10043CV) with B-27 supplement without antioxidants (2%) (Gibco, #10889038). During RI, cells were incubated with CM(+) medium alone or supplemented with decellularized ECM (1x or 5x concentration). For functional recovery, spontaneous beatings were recorded at 1h, 3h, 6h and 12h RI. At 3h RI, cellular ROS was measured and at 12h RI apoptosis-related proteome was profiled (R&D Systems, #ARY009), and survival rate was measured via live/dead staining (Abcam, #ab115347).

Aged cells were treated with decellularized ECM for 10 days and control groups were maintained in CM(+) media throughout the experiment. Cells were screened for their relative cytokine content and gene expressions before and after ECM treatment. After treatment, spontaneous beating of the cells as well as mitochondrial health (ThermoFisher, MitoProbe JC-1, #M34152) and cellular ROS (Abcam, Cellular ROS Assay #ab186029) were assessed.

### 5.6. RNA Isolation

Cells were rinsed with PBS, collected with trypsin, and stored in a −80°C freezer for future RNA isolation. For RNA isolation, frozen cells were thawed and centrifuged to remove the freezing media. The pellet was then processed following the RNeasy Mini Kit (Qiagen, #74104) protocol. Briefly, cells were disrupted using the lysis buffer and same volume ethanol added to the lysate. The sample is then applied to the RNeasy mini spin column, collected on the membrane, and finally RNA was eluted in RNAse-free water. RNA purity was confirmed, concentration was measured using a Nanodrop 2000 spectrophotometer, and samples were sent to the core facility at Ohio State University.

### 5.7. Gene Expression Analysis

mRNA levels were quantified using NanoString Technology. An nCounter custom codeset was designed for the identification of genes of interest related to iCM maturity, function and apoptosis with a total of 64 genes including 5 housekeeping genes (*B2M*, *EEF1A1, GAPDH, RNPS1,* and *SRP14*), selected based on a publication [43]. RNA inputs of 100 ng were used for hybridization and placed on a cartridge for the NanoString reader. The output files (RCC files) were loaded into nSolver Analysis software. Data was run for quality control and background normalization, then genes of interest were normalized to the housekeeping genes. For visualization purposes, heatmap of log2FC for the differentially expressed genes was generated using the nSolver analysis software v4.0. Complete linkage hierarchical clustering method with Euclidean distance was used to cluster the human tissues and iCMs.

Gene Ontology (GO) Analysis was performed on the obtained relative expression data. Comprehensive analysis was performed using an online database via EnrichR for biological processes and enrichment analysis[44]. Data was extracted from the output dataset and graphed manually using the p-value provided. Proteomic interactions of the same relative expression data were also classified through KEGG-based proteomapping software[45] and are presented as obtained.

### 5.8. Proteome Analysis

The relative cytokine content of the aged iCMs before and after ECM treatment and of the decellularized human heart ECM were obtained using the Human XL Cytokine Array Kit (ARY022B, R&D Systems). Briefly, cell lysates were obtained by disrupting the cells using the lysis buffer supplemented with protease inhibitor cocktail and tissue extracts were obtained by 1% Triton-X incubation. The relative expression levels of apoptosis related proteins of post I/RI iCMs (n=3) were analyzed using the Proteome Profiler Human Apoptosis Array (R&D Systems, #ARY009), according to the manufacturer’s instructions.

For all, protein concentrations were normalized before starting the arrays and incubated overnight with pre-blocked membranes at 4°C. At the end, the unbound proteins were rinsed away, the membranes were incubated with the Streptavidin-HRP, then developed using chemiluminescent detection reagent mixture. For quantification, the background was removed, and the pixel density of each spot was measured using ImageJ.

### 5.9. Beating Analysis

A block-matching algorithm was performed using MATLAB as described previously[6]. Briefly, spontaneous beating of iCMs were recorded (n=3-4 ROI per sample) and the beating frequency, average beating velocity, average maximum displacement, and beat area (%) were calculated. To analyze, measurements were normalized to pre-treatment values.

### 5.10. Statistical Analysis

The mean ± standard deviation (SD) was reported for all replicates. One-way ANOVA with post-hoc Tukey’s test was used to assess the statistically significant differences using GraphPad Prism version 8. All *p* values reported were two-tailed, and *p* < 0.05 was considered statistically significant. Sample size (n) ≥ 3 for individual experiments.

## Acknowledgements

This work was supported by the National Science Foundation CBET grant number [1651385] and [NSF-1805157]. We would also like to thank Dr. Keith L March and Indiana Donor Network for facilitating the human tissue acquisition and Dr. Bradley Ellis for transportation.

## Data availability statement

The raw/processed data required to reproduce these findings can be shared upon reasonable request.

## Ethics Statement

Deidentified human hearts from donors that were deemed unsuitable for transplantation and donated to research were collected through the Indiana Donor Network under the Institutional Review Board (IRB) approval for deceased donor tissue recovery. All human tissue collection conformed to the Declaration of Helsinki.

